# A rational approach to identifying effective combined anticoronaviral therapies against feline coronavirus

**DOI:** 10.1101/2020.07.09.195016

**Authors:** S.E. Cook, H. Vogel, D. Castillo, M. Olsen, N. Pedersen, B. G. Murphy

## Abstract

Feline infectious peritonitis (FIP), caused by a genetic mutant of feline enteric coronavirus known as FIPV, is a highly fatal disease of cats with no currently available vaccine or FDA-approved cure. Dissemination of FIPV in affected cats results in a range of clinical signs including cavitary effusions, anorexia, fever and lesions of pyogranulomatous vasculitis and peri-vasculitis with or without central nervous system and/or ocular involvement. There is a critical need for effective and approved antiviral therapies against coronaviruses including FIPV and zoonotic coronaviruses such as SARS-CoV-2, the cause of COVID-19. With regards to SARS-CoV-2, preliminary evidence suggests that there may be potential clinical and pathological overlap with feline coronaviral disease including enteric and neurological involvement in some cases. We have screened 89 putative antiviral compounds and have identified 25 compounds with antiviral activity against FIPV, representing a variety of drug classes and mechanisms of antiviral action. Based upon successful combination treatment strategies for human patients with HIV or hepatitis C virus infections, we have identified combinations of drugs targeting different steps of the FIPV life cycle resulting in synergistic antiviral effect. Translationally, we suggest that a combined anticoronaviral therapy (cACT) with multiple mechanisms of action and penetration of all potential anatomic sites of viral infection should be applied towards other challenging to treat coronaviruses, like SARS-CoV-2.

**Author summary:** We have screened 89 compounds *in vitro* for antiviral activity against FIPV. The putative antiviral activity of these compounds was either purported to be a direct effect on viral proteins involved in viral replication or an indirect inhibitory effect on normal cellular pathways usurped by FIPV to aid viral replication. Twenty-five of these compounds were found to have significant antiviral activity. Certain combinations of these compounds were determined to be superior to monotherapy alone.

## Introduction

Feline Infectious Peritonitis (FIP) is a highly fatal disease of cats with no effective vaccine or FDA-approved treatment. Although the pathogenesis has not been fully elucidated, FIP is generally understood to develop as the result of specific mutations in the viral genome of the minimally pathogenic and ubiquitous feline enteric coronavirus (FECV), creating the virulent FIP virus (FIPV)[1–3]. These FECV mutations result in a virus-host cell tropism switch from intestinal enterocytes to peritoneal-type macrophages. Productive macrophage infection by FIPV, targeted widespread anatomic dissemination, and immune-mediated perivasculitis results in the highly fatal systemic inflammatory disease, FIP[4]. As a result of viral dissemination, FIP may present with clinical signs reflecting inflammation in a variety of anatomic sites potentially including the abdominal cavity and viscera, thoracic cavity, central nervous system and/or eye[5–8]. FIP remains a devastating viral disease of cats due to its high mortality rate, challenges in establishing a precise etiologic diagnosis, and the current lack of available and effective treatment options[7, 9]. The development of an effective vaccine for FIP has been complicated by the role of antibody-dependent enhancement (ADE) in FIP disease pathogenesis, where the presence of non-neutralizing anti-coronaviral antibodies have been shown to exacerbate FIP disease[10–12].

Mammalian coronaviruses infect and generally cause disease in either the intestinal tract or respiratory system of their infected vertebrate hosts[13]. However, FIP often manifests as a multisystemic inflammatory disease syndrome as a result of the widespread dissemination of FIPV-infected macrophages. The recent pandemic emergence of SARS-CoV-2 has resulted in variable disease syndromes in infected human patients, collectively referred to as COVID-19. Although SARS CoV-2 has an apparent tropism for respiratory epithelium resulting in interstitial pneumonia, recent evidence indicates that COVID-19 can also present as alimentary disease, manifesting clinically as diarrhea[14, 15]. Tropism for these tissues reflects membrane expression of the ACE2 protein, the cellular target of SARS-CoV-2[16]. SARS-CoV-2 further appears to be capable of infecting and causing inflammatory disease in tissues external to intestinal tract and respiratory systems, including the brain, eye, reproductive tract, and cardiac myocardium[17–21]. Brainstem neuroinvasion and subsequent encephalitis by SARS CoV-2 may contribute to respiratory failure in COVID-19 patients[20, 22]. Experimentally, SARS CoV-2 is also able to establish a productive infection in cats[23]. Therefore, FIPV infection of cats and SARS CoV-2 infection of human patients bear more resemblance than may have been initially perceived.

There is an immediate and critical need for available and effective antiviral therapies to treat these coronaviral diseases. FIPV-infected cats could serve as a translational model and provide useful guidance for SARS CoV-2 patients with COVID-19. Recent antiviral clinical trials in both experimentally and naturally FIPV-infected cats have shown promise in treating and curing FIP through the use of GS-441524, a nucleoside analog and metabolite of the prodrug Remdesivir (Gilead Sciences), or GC-376, a 3C-like protease inhibitor of FIPV (Anavive)[24–26]. The GS-441524 prodrug Remdesivir has recently shown promise in treating human patients infected with SARS CoV-2[27, 28]. Despite these recent clinical successes, these antiviral compounds have yet to be approved and are currently unavailable for clinical veterinary use in cats with FIP.

The identification and development of effective antiviral therapies can be both costly and time consuming. The targeted screening and repurposing of drugs previously FDA-approved or approved for research use can play an efficient role in drug discovery. Using a slate of putative antiviral compounds selected based on their proven efficacy in treating other RNA-viruses, we identified a subset of compounds with strong anti-FIPV activity and characterized their safety and efficacy profiles *in vitro*. Based upon the great success of combined anti-retroviral therapy (cART) against HIV-1 and combination therapies against hepatitis C virus[29], we have developed methods to identify effective combinational therapies against FIPV. Initial monotherapies against HIV-1, such as azidothymidine (AZT), often resulted in viral escape mutations. The use of multiple antiviral compounds concurrently appears to block this adaptive viral evolutionary mechanism as HIV-1 evolution is effectively halted by modern day cART[30]. The success of cART is thought to be the result of pharmacologically targeting the virus lifecycle at multiple stages simultaneously, collectively achieving a synergistic antiviral effect[31].

Given the impressive success of cART, it would seem feasible that concurrently targeting FIPV at different steps of the virus lifecycle with a combined anti-coronaviral therapy (cACT) may offer a greater level of sustained and more complete success than has been achieved with monotherapies alone. The inclusion of an antiviral agent in cACT capable of penetrating the blood-brain-barrier (BBB) and eye and achieving pharmacologically relevant tissue concentrations may facilitate system-wide FIPV eradication. Here we describe a set of *in vitro* assays facilitating the rapid screening and identification of effective anti-coronaviral compounds. Efficacious antiviral agents with different mechanisms of action and predicted body distribution were combined into cACT and tested for compound synergy. We hypothesized that the combinatorial use of two or more effective antiviral monotherapies with differing mechanisms of action will facilitate the identification of synergistic combinations providing superior anti-coronaviral efficacy compared to their use as sole agents. The identification of successful cACT may also provide guidance towards the treatment of other emerging viral diseases, such as SARS-CoV-2.

## Results

### Compound Screening

In order to identify compounds with anti-FIPV activity, a compilation of 89 compounds (Supplementary Table 1) from differing drug classes and with a variety of putative mechanisms of action were screened for anti-coronaviral activity in *in vitro* assays. Compounds screened included nucleoside polymerase inhibitors (NPIs), non-nucleoside polymerase inhibitors (NNPIs), protease inhibitors (PIs), NS5A inhibitors, a set of novel anti-helicase chemical “fragments”, and a set of compounds with undetermined mechanisms of action. From this group of 89 compounds, a total of 25 different compounds were determined to possess antiviral activity against FIPV including NPIs, PIs, NS5A inhibitors and two compounds with undetermined mechanisms of action (termed “other”, Fig 1). These successful antiviral compounds included toremifene citrate, daclatasvir, elbasvir, lopinavir, ritonavir, nelfinavir mesylate, K777/K11777, grazoprevir, amodiaquine, EIDD 1931, EIDD 2801, and GS-441524 sourced from three different China-based manufacturers (Table 1, Fig. 2). We tested several nucleoside analog compounds provided by Gilead Sciences structurally related to the nucleoside analogs GS-441524 and Remdesivir for their antiviral properties and found several with potential (included in the above reported 25 identified compounds) but did not pursue these agents further. As a result, the total number of antiviral agents carried forward for further analyses was 13. This total includes the previously identified 3-C protease inhibitor, GC-376 (Anavive).

**Table 1.** EC50 of compounds with anti-FIPV activity.

| Compound Name | Drug Category | EC50 (μM) |
| --- | --- | --- |
| GC376 | PI | 0.04 |
| EIDD 1931 | NPI | 0.09 |
| Elbasvir | NS5A Inhibitor | 0.16 |
| EIDD 2801 | NPI | 0.4 |
| GS-441524<sup>†</sup> | NPI | 0.66 |
| K777/K11777 | PI | 0.67 |
| Toremifene citrate | Other* | 5 |
| Amodiaquine | Other** | 6.5 |
| Lopinavir | PI | 8.08 |
| Ritonavir | PI | 8.7 |
| Grazoprevir | PI | 12.13 |
| Nelfinavir mesylate | PI | 13.47 |
| PI = Protease Inhibitor; NPI = Nucleoside Polymerase Inhibitor |  |  |
| <sup>†</sup> MedChem Express, HY-103586 |  |  |
| *Selective Estrogen Receptor Modulator |  |  |
| **4-Aminoquinoline |  |  |

**Figure 1.**
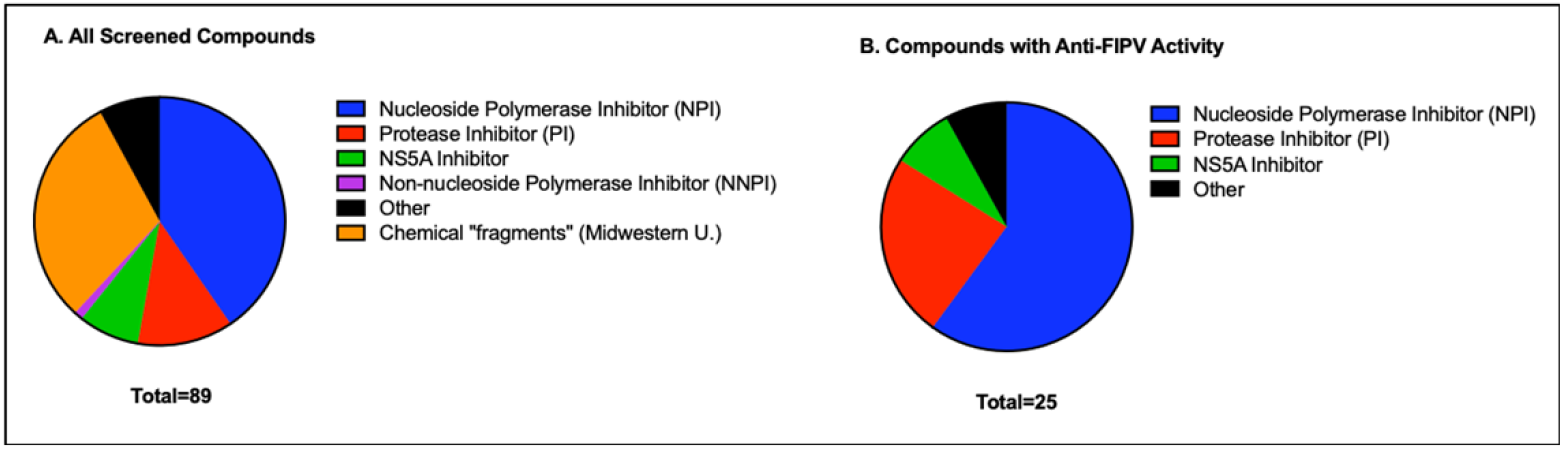
Compounds screened by mechanism of action. (A) Pie graph depiction of all compounds screened. (B) Compounds identified during screening to possess anti-FIPV activity *in vitro*.

**Figure 2.**
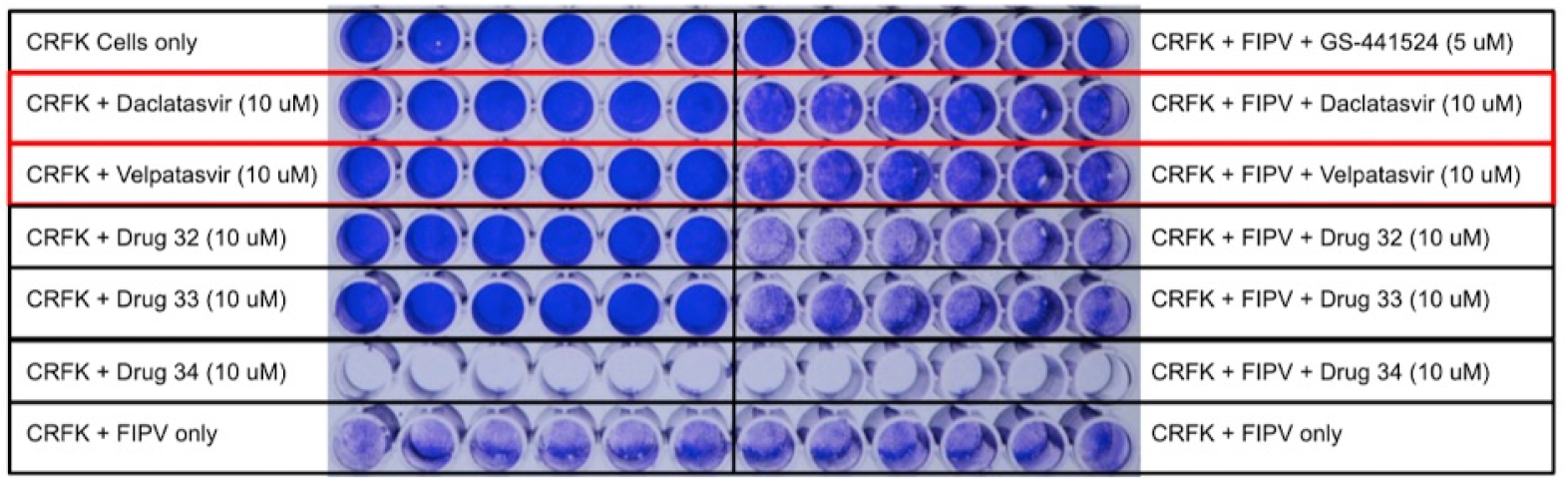
Example screening plate using crystal violet staining to identify anti-FIPV activity at 10 μM. The top left row are control wells with CRFK cells only and no drug or FIPV. The top right row is a positive control utilizing GS-441524 with known complete protection of CRFK cells against FIPV-induced cell death. The entire bottom row of wells represents CRFK cells infected with FIPV and no drug treatment. The remaining rows are screening wells with the left half assessing for cytotoxicity at 10 μM (no FIPV infection) and the right half assessing for anti-FIPV activity at 10 μM for any given compound. Loss of staining indicates cell loss. Daclatasvir and velpatasvir demonstrated anti-FIPV activity evidenced by increased crystal violet staining (relatively intact cell monolayers) relative to FIPV-only control wells (bottom row of plate). Drugs 32, 33, and 34 demonstrated absent to minimal antiviral effect, with drug 34 (Ravidasvir) also demonstrating cytotoxicity at 10 μM based on the dramatic well clearing seen on the left half of the plate without FIPV.

### Determining antiviral efficacy

The antiviral efficacy (EC50) was determined for 10 antiviral compounds. For these compounds, the EC50 ranged from 0.04 μM to 13.47 μM (Table 1, Fig. 3). One of the antiviral agents, Daclatasvir, demonstrated unacceptable cytotoxicity at 20 μM and was removed from further testing. GS-441524 sourced from China (MedChemExpress, HY-103586) was shown to have a comparable EC50 relative to previously published values for GS-441524 sourced from Gilead Sciences[25].

**Figure 3.**
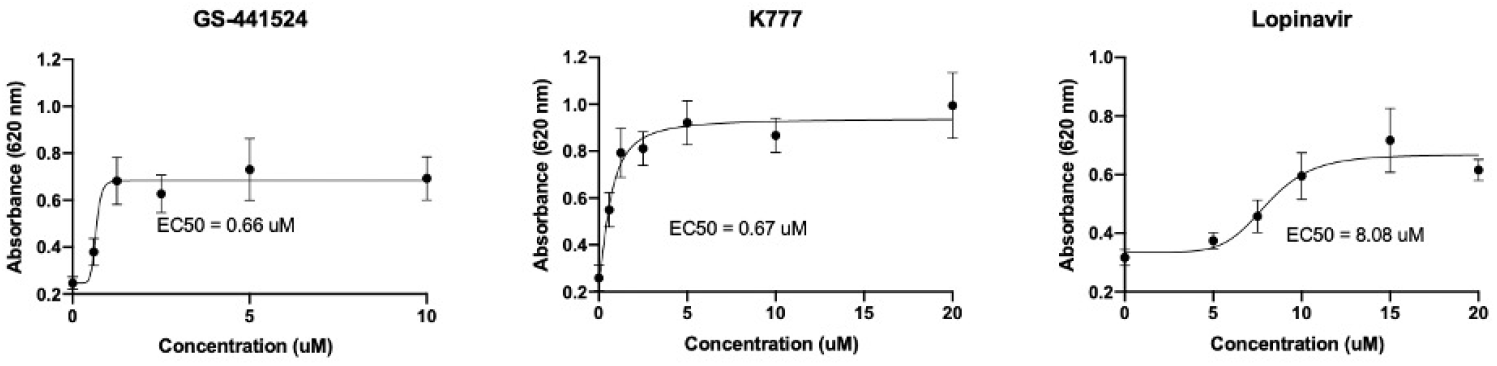
Representative examples of EC50 nonlinear regression analyses for compounds with anti-FIPV activity. Serial dilutions of each compound with anti-FIPV activity were performed to identify the half maximal effective concentration (EC50). GS-441524 results shown here represents the compound sourced from MedChemExpress.

### Cytotoxicity Safety Profiles

Cytotoxicity Safety Profiles (CSP) were determined for ten different antiviral compounds in CRFK cells. At 5 μM, seven of the tested compounds demonstrated essentially no cytotoxicity while two of the antivirals, amodiaquine and toremifene had 11 and 12% cytotoxicity, respectively (Fig. 4; Table 2). The 50% cytotoxic concentration (CC50) for GC376 has been reported previously as > 150 μM [32]. Interestingly, based on the Promega CellTox™ Green Cytotoxicity Assay, the cytotoxicity of both EIDD compounds was essentially undetectable up to 100 μM. However, visual inspection of the EIDD treated wells just prior to fluorescent dye application and plate readings revealed differences in cell morphology (cytopathic effect) between untreated CRFK cells and treated cells. The untreated CRFK cells were characterized by adherent spindled morphology in a single monolayer, while the EIDD-treated wells demonstrated a clear decrease in confluency by comparison with variable cell morphology including rounding up of cells (cytopathic effect). The inconsistency between subjective visual assessment of EIDD-treated wells and the fluorescence assay is enigmatic. It is possible that the overall decreased cell number in EIDD-treated wells resulted in loss and degradation of nucleic acid necessary for fluorescence binding and detection in the CellTox assay.

**Table 2.** Percent cytotoxicity by compound and concentration.

| Compound Name | Drug Category | 5 uM | 10 uM | 25 uM | 50 uM | 100 uM |
| --- | --- | --- | --- | --- | --- | --- |
| Elbasvir | NS5A Inhibitor | 0.67 | 0.46 | 1.4 | 2 | 4.9 |
| K777/K11777 | PI | 0.61 | 0.29 | 2.3 | 9 | 16 |
| Toremifene citrate | Other* | 12 | 22 | 23 | 35 | 39 |
| Amodiaquine | Other** | 11 | 12 | 19 | 23 | 26 |
| Lopinavir | PI | 0.67 | 15 | 13 | 16 | 18 |
| Ritonavir | PI | 0.49 | 0.84 | 2 | 33 | 26 |
| Grazoprevir | PI | 0.55 | 0 | 0.81 | 3.9 | 19 |
| EIDD 1931 | NPI | 1.2 | 0.94 | 0.6 | 0.78 | 2.8 |
| EIDD 2801 | NPI | 0 | 0.95 | 0.63 | 1.7 | 0.8 |
| GS-441524<sup>†</sup> | NPI | 0.21 | 0 | 0 | 0 | 0 |

PI = Protease Inhibitor; NPI = Nucleoside Polymerase Inhibitor \*Selective Estrogen Receptor Modulator \*\*4-Aminoquinoline <sup>†</sup>NMPharmTech

**Figure 4.**
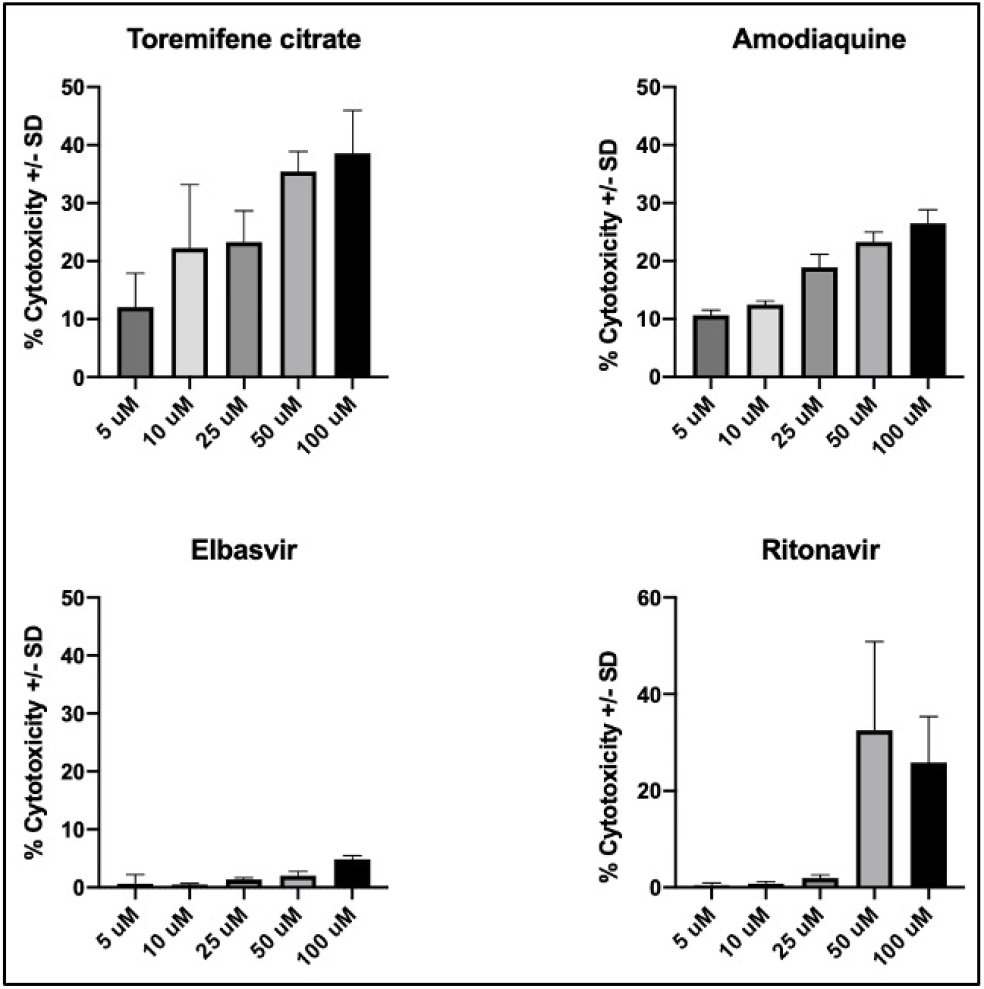
Representative cytotoxicity profiles. Percent cytotoxicity bar graphs +/− standard deviation (SD) for four compounds with anti-FIPV activity. Percent cytotoxicity values were determined by normalizing cytotoxicity to the positive toxicity control wells (set to 100% cytotoxicity) and untreated CRFK cells (set to 0% baseline cytotoxicity).

### Quantification of compound inhibition of viral RNA production with monotherapy

A real-time RT PCR assay was utilized to measure each antiviral compounds’ ability to inhibit coronaviral replication as monotherapy (viral RNA knock-down assay). Compounds demonstrating the greatest inhibition of FIPV RNA production were GC376, a 3C-like coronavirus protease inhibitor, GS-441524, EIDD-1931 and EIDD-2801, the latter three all being nucleoside analogs (Fig. 5, Table 3). Those with the least inhibitory effect on viral RNA production include elbasvir, nelfinavir, and ritonavir. Ritonavir, a protease inhibitor, is used in combination with lopinavir to treat HIV-1 infection (Kaletra, AbbVie). Lopinavir monotherapy has poor oral bioavailability in people, however, when used in combination, Ritonavir has been demonstrated to markedly improve lopinavir’s plasma concentration[33]. Therefore, despite the relatively minimal FIPV inhibition identified with ritonavir as monotherapy, this compound was carried forward for additional testing, including combined anticoronaviral assessment.

**Table 3.** Fold reduction in viral RNA copy number for anti-FIPV compounds.

| **Monotherapy Fold Reduction in FIPV RNA** |  |
| --- | --- |
| **Compound** | **Fold Reduction in Viral Titer** |
| GC376 (20 uM) | 25,000 |
| GC376 | 7,300 |
| GS-441524 (NMPharmTech) | 5,280 |
| EIDD-1931 | 3,700 |
| GS-441524 (MedChemExpress) | 3,500 |
| EIDD-2801 | 2,110 |
| Lopinavir | 309 |
| Toremifene | 10 |
| K777 | 7 |
| Grazoprevir | 5 |
| Amodiaquine | 4 |
| Elbasvir | 2 |
| Nelfinavir mesylate | 1 |
| Ritonavir | 1 |

\*Unless otherwise indicated, all compounds utilized at 10 uM.

**Figure 5.**
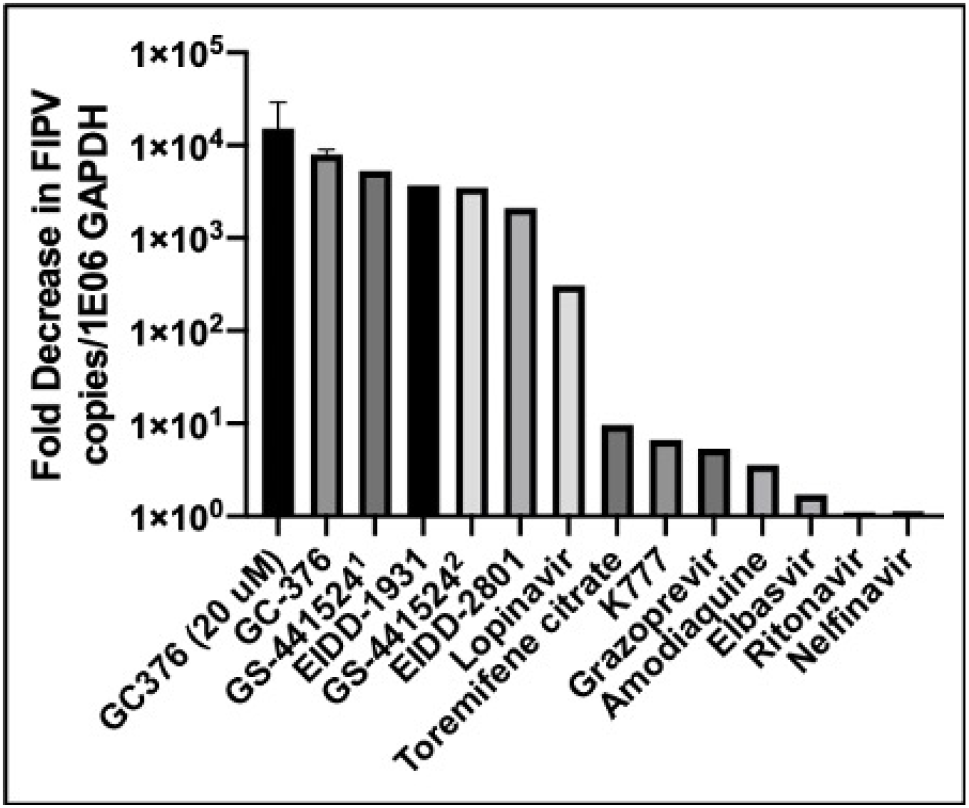
Fold decrease in FIPV RNA copy number utilizing antiviral compounds as monotherapy. FIPV-infected CRFK cells were incubated for 24 hours with compounds identified to possess anti-FIPV activity. Viral copy number was subsequently determined via RT-qPCR and normalized to feline GAPDH copy number to determined fold decrease effect for each compound. All compounds were tested at 10 μM unless otherwise specified. All experimental treatments were performed in triplicate wells and fold decrease calculated by dividing the average experimental, normalized FIPV copy number by the average normalized FIPV copy number determined for untreated, FIPV-infected wells. ^1^*GS-441524 sourced from NMPharmTech (China)*. ^2^*GS-441524 sourced from MedChemExpress (China)*.

### Quantification of compound inhibition of viral RNA production with cACT

In order to identify drug combinations with synergistic antiviral activity over monotherapy, combinations of two or more drug compounds were selected based upon (i) established combinations utilized for other viral infections like HIV-1 and HCV, (ii) drugs with different mechanisms of action, (iii) potential variations in systemic distribution of the compound (e.g. chemical class-based ability to penetrate the blood-brain or blood-ocular barriers), and (iv) minimal cytotoxicity (based on the CSP). For each cACT, any resulting decrease in FIPV copy number over the calculated additive effect for each drug used as monotherapy was considered to be synergistic (Table 4). The combination of antiviral agents with the greatest total fold reduction in viral RNA as well as greatest synergistic effect was determined to be GC376 and amodiaquine with a 76-fold decrease in viral RNA over the additive effect (Fig 6). This particular synergistic combination was one of the more interesting results given that amodiaquine alone demonstrated limited inhibition of FIPV viral RNA copies determined by qRT-PCR.

**Table 4.** Fold reduction in FIPV viral RNA copy number using combinatorial therapy (cACT). The expected additive effect reflects the sum of the fold reduction in viral RNA based on each agent used as monotherapy (Table 3).

| Compound 1 | Compound 2 | Compound 3 | Fold Reduction in Viral Titre | Additive | Fold over additive |
| --- | --- | --- | --- | --- | --- |
| GC376 (20 μM) | Amodiaquine | -- | 1,897,000 | 25004 | 76 |
| GC376 (20 μM) | Amodiaquine | Toremifene | 256,000 | 25014 | 10 |
| GC376 (20 μM) | K777 | -- | 248,000 | 25007 | 10 |
| GC376 (20 μM) | Toremifene | -- | 128,000 | 25010 | 5.1 |
| GC376 (20 μM) | Nelfinavir mesylate | -- | 91,100 | 25001 | 3.6 |
| Elbasvir (5 μM) | Lopinavir | -- | 14,000 | 311 | 45 |
| Elbasvir (5 μM) | GC376 (20 μM) | -- | 12,600 | 25002 | 0.50 |
| GC376 (10 μM) | Amodiaquine | -- | 11,700 | 7304 | 1.6 |
| GC376 (10 μM) | Grazoprevir | -- | 8,290 | 7305 | 1.1 |
| GC376 (20 μM) | GS-441524 (NMPharmTech) | -- | 8,260 | 30280 | 0.27 |
| GC376 (10 μM) | Amodiaquine | Elbasvir (5 μM) | 7,570 | 7306 | 1.0 |
| GC376 (10 μM) | GS-441524 (MedChem) | -- | 6,910 | 10800 | 0.64 |
| GC376 (20 μM) | Ritonavir | -- | 6,560 | 25001 | 0.26 |
| K777 | Lopinavir | -- | 5,530 | 316 | 18 |
| GC376 (10 μM) | GS-441524 (NMPharmTech) | -- | 4,340 | 12580 | 0.34 |
| GC376 (20 μM) | Lopinavir | -- | 3,400 | 25309 | 0.13 |
| Lopinavir | Ritonavir | -- | 3,130 | 310 | 10 |
| Lopinavir | Ritonavir | Toremifene | 1,740 | 320 | 5.4 |
| GC376 (10 μM) | EIDD-2801 | -- | 255 | 9410 | 0.03 |
| K777 | Toremifene | -- | 25 | 17 | 1.5 |

\*Unless otherwise indicated, all compounds utilized at 10 μM.

**Figure 6.**
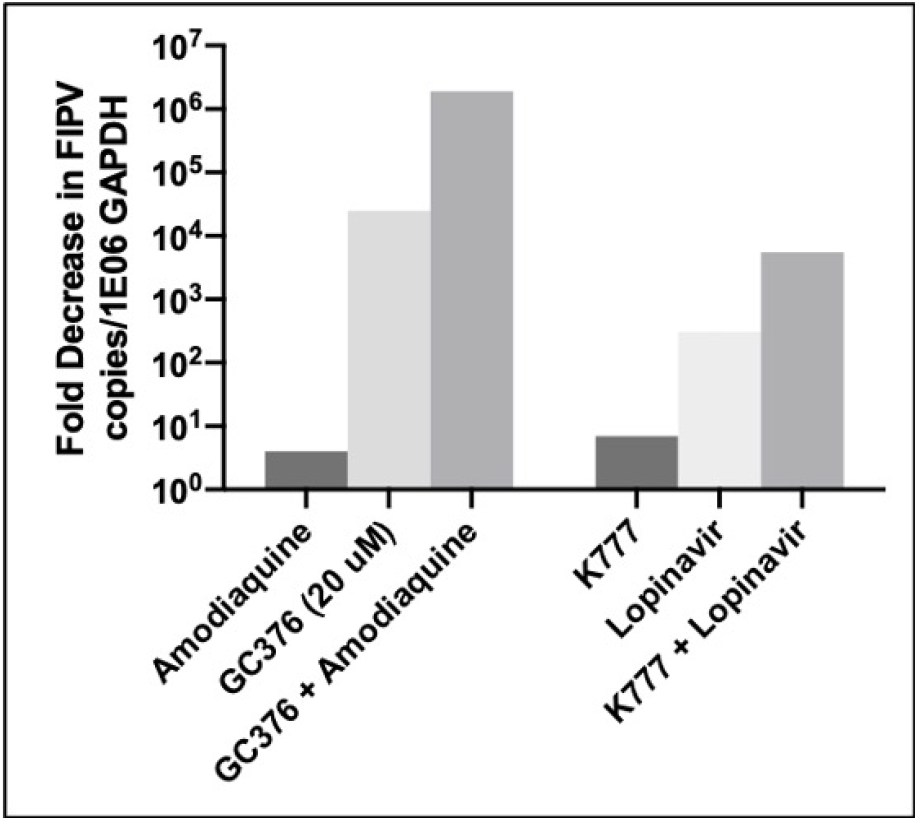
Select examples of fold reductions of FIPV RNA copy number using combinatorial therapy (cACT). Bars represent the mean fold decrease of three treated wells of CRFK cells compared to the mean fold decrease of three untreated, FIPV-infected wells. All compounds tested at 10 μM unless otherwise indicated.

Due to pronounced anti-FIPV activity of GC-376 as well as its potential availability for moving forward into *in vivo* pharmacokinetic studies, clinical trials, and hopeful use in an established cACT application, this compound was focused on in a series of mono- and combined therapy “viral RNA knock-down” assays (Fig 7). Overall, GC376 demonstrated superior anti-FIPV activity at 20 μM both as monotherapy and in combinatorial therapies *in vitro*. The most marked reduction in FIPV RNA occurred with combinations of GC376 at 20 μM, in particular with amodiaquine at 10 μM. The experiment combining GC376 with amodiaquine was repeated and both results are reported in Fig 7C for comparison.

**Fig 7.**
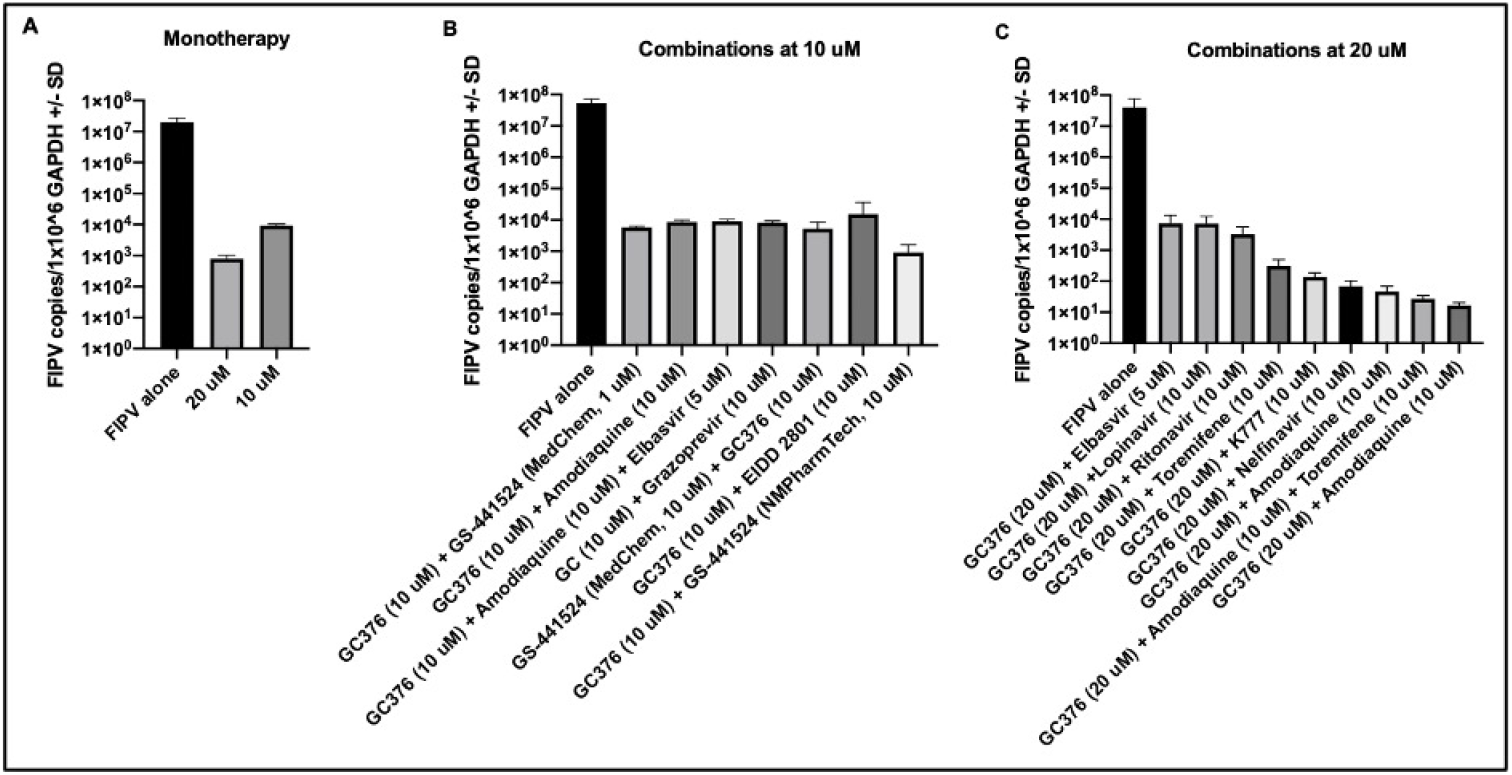
GC376 antiviral activity as monotherapy and in combinations at 10- and 20 μM. (A) Reduction of FIPV RNA quantified by RT-qPCR using GC376 as monotherapy at 10- and 20 μM. There is a significant difference between these two concentrations with 20 μM superior to 10 μM. (unpaired t-test; *p*<0.0001). (B) Combination *in vitro* therapy using GC376 at 10 μM. (C) Combination *in vitro* therapy against FIPV using GC376 at 20 μM.

## Discussion

As there is no currently effective vaccine for FIP, there is a strong clinical and world-wide need for effective antiviral treatment options for FIPV-infected cats. Here we describe the screening of 89 compounds, resulting in the identification of 25 antiviral agents with antiviral efficacy and strong safety profiles against the feline coronavirus, FIPV. We also identified combinations of antiviral agents (cACT) that resulted in superior efficacy, or synergism, over monotherapy alone. Particularly interesting was a finding relating to the use of elbasvir, which repeatedly demonstrated excellent CRFK protection from FIPV-induced CPE at less than 1 μM based on multiple plaquing assays (EC50 of 0.16 μM). However, essentially no difference was detected in the viral RNA copy number between infected cells treated with or without elbasvir. Further visual analysis of FIPV infected CRFK cells that were treated with elbasvir revealed an atypical cell morphology relative to the uninfected cells characterized by variable swelling, rounding of cells and scattered partial cell detachment (cytopathic effect). These “atypical cells” sparsely detached from the culture plate and as a result, the absorbance values acquired in the plaquing assay were comparable to uninfected control wells. This dichotomous result between the plaque assay and viral RNA knock-down assay suggests that the antiviral effect of elbasvir may be downstream of viral replication, and as a result, elbasvir may not protect cells from the accumulation of viral RNA. Elbasvir has been utilized for the treatment of hepatitis C virus (HCV) infected patients and is thought to target the HCV NS5A protein preventing replication and also virion assembly[34]. Although an NS5A homolog is not identified for FIPV, it is possible that elbasvir exerts a similar antiviral effect by preventing FIPV virion assembly without blocking viral RNA synthesis in CRFK cells. Additional assessment of FIPV-infected and treated CRFK cells using transmission electron microscopy may provide clarity of elbasvir’s effect on CRFK protection from FIPV-associated cellular injury and death.

The coadministration of ritonavir with lopinavir has been shown to markedly enhance the plasma concentration of lopinavir in rats, dogs, and humans[33]. Ritonavir is a potent inhibitor of CYP3A, which is the primary enzyme responsible for the metabolism of protease inhibitors, hence, its coadministration with other protease inhibitors results in enhanced systemic concentrations of the co-administered protease inhibitor, such as lopinavir[38, 39]. Ritonavir-associated augmentation of the antiviral effect of lopinavir was relatively minimal in the viral RNA knock-down assays with only a 10-fold FIPV inhibition over additive effect. This may be the result of an *in vitro* artifact of testing in a feline renal cell line (i.e. CRFK cells) lacking the CYP3 enzyme, an enzyme typically seen in sites of high protease inhibitor first-pass metabolism (i.e., enterocytes and hepatocytes)[39]. These results suggest that *in vitro* assays alone may not fully predict the effect of antiviral agents in FIPV infected cats *in vivo*.

Grazoprevir, an NS3/4 serine protease inhibitor, has been used in combination with elbasvir, an NS5A inhibitor, to treat patients infected with HCV (Zepatier, Merck)[40]. Here we have demonstrated that grazoprevir possesses anti-FIPV activity when used as monotherapy. The cysteine protease inhibitor K777/K11777 has been investigated for its ability to block coronavirus entry (MERS-CoV and SARS-CoV-1) and ebolavirus and was found to fully inhibit coronavirus infection but only in target cell lines lacking virus-activating serine proteases[41]. For other cell lines, K777 inhibited coronavirus cell entry when combined with a serine protease inhibitor[41]. It is possible that K777’s limited inhibition of FIPV RNA production could be augmented if combined with a serine protease inhibitor.

Amodiaquine is an antimalarial drug and member of the 4-aminoquinoline drug class. Amodiaquine, along with related 4-aminoquinolines such as chloroquine and hydroxychloroquine, was originally developed for the treatment of malaria[42], and like chloroquine and hydroxychloroquine, possesses widespread anatomical volume of distribution, including eyes and brain[43–49]. Antiviral compound penetrance into the CNS and/or ocular compartments is particularly relevant in the case of FIPV-infected cats with neurologic and/or ocular involvement. While several investigations have defined the antiviral properties of chloroquine and hydroxychloroquine[28, 50, 51], the antiviral activity of amodiaquine has also been investigated with the identification of antiviral activity against dengue virus, Ebola virus, and severe fever with thrombocytopenia syndrome (SFTS) virus[52–55]. The mechanism of action of amodiaquine may involve an increase of cytoplasmic lysosomal and/or endosomal pH, preventing the release of viable virions into the cytoplasm[56]. Given its unique drug class status and purported ability to cross the blood-brain-barrier[57] amodiaquine holds promise as a component of combinatorial therapy for the treatment of neurologic and/or ocular FIP.

Toremifene citrate, a selective estrogen receptor modulator (SERM), has been used for the treatment of metastatic breast cancer in human patients. More recently, toremifene has been evaluated for its antiviral properties and has demonstrated anticoronaviral activity against the zoonotic coronaviruses Middle Eastern Respiratory Syndrome Coronavirus (MERS-CoV) and SARS-CoV-1[58]. Toremifene has also demonstrated activity against the Ebola virus (EBOV)[59, 60]. While the exact mechanism of antiviral action has not been defined, toremifene’s antiviral action against EBOV appears to be due to destabilization of the EBOV glycoprotein[59].

Interestingly, GC376 demonstrated perplexing differences between dosing at 10 μM versus 20 μM in combination therapy. When used in combination at 10 μM with other compounds, there was absent synergism and in some cases a decrease in antiviral effect with fold over additive values ranging from 0.03 to 1.6 (Table 4). When used at 20 μM there were still instances where combining with another compound resulted in a decreased antiviral effect compared to GC376 used as monotherapy at 20 μM. However, there was much more variation with combinations at 20 μM with fold over additive values ranging from 0.13 to 76 (Table 4). One particular example is the contrast between GC376 at 20 μM compared with GC376 at 10 μM combined with amodiaquine at 10 μM. The former resulted in the greatest viral RNA inhibition as well as the greatest fold over additive (synergistic) effect, while the latter nearly lost synergism with a fold over additive value of 1.6.

The identification of effective antiviral strategies for treating FIPV-infected cats holds translational implications for the ongoing SARS-CoV-2 pandemic. FIPV-infection in cats is reminiscent of coronavirus infection in ferrets [61, 62] and has been compared to the pathogenesis of other chronic, macrophage-dependent diseases such as tuberculosis[63]. As the clinical and pathogenic details of SARS-CoV-2 infection in people continues to emerge, there appears to be some overlap with FIPV in anatomic distribution, clinical manifestation, and likely, response to certain antiviral therapies. In cats, the feline enteric coronavirus biotype (FECV) is restricted to the alimentary tract as a result of an enterocyte tropism. Clinical signs in cats infected with FECV range from mild gastrointestinal disease (diarrhea) to absent. The mutated feline coronavirus biotype FIPV acquires a macrophage tropism and preferentially targets serosal surfaces of the abdominal and thoracic cavities with a subset of cats demonstrating CNS or ocular involvement[7]. Similarly, for COVID-19 patients there are reports of diarrhea and a subset of patients with CNS-involvement[14]. Although the cellular receptor for SARS-CoV-2 has been identified as ACE2[64], the cellular receptor for serotype I FIPV has yet to be determined. The cellular receptor for the less clinically relevant FIPV serotype II has been identified as feline aminopeptidase peptidase (fAPN)[4]. A study utilizing RNAseq to evaluate gene expression profiles of ascites cells obtained from cats with FIP did not identify expression of ACE2, suggesting that ACE2 is unlikely to be the serotype I FIPV receptor[63]. More investigation into the identity of the serotype I FIPV receptor is warranted.

Past clinical successes using GS-441524 or GC-376 in cats with experimental and naturally occurring FIP demonstrate that an effective cure for FIP is possible, however, challenges in treating non-effusive (granulomatous), neurological and ocular FIP remain. The 3C-like protease inhibitor, GC-376, appears to be relatively effective in the treatment of effusive FIPV infection confined to body cavities but may be less effective in treating the neurological or ocular forms of the disease[24]. These differing outcomes may be the result of inefficient penetration of the blood-brain and blood-eye barriers, making GC-376 a promising candidate for combined therapy with a CNS-penetrating antiviral drug.

## Materials and Methods

### FIPV inoculum for in vitro experiments

Crandell-Reese feline kidney cells (CRFK, ATCC) were cultured in T150 flasks (Corning), inoculated with serotype II FIPV (WSU-79-1146, GenBank DQ010921) and propagated in 50 mL of Dulbecco’s Modified Eagle’s Medium (DMEM) with 4.5g/L glucose (Corning) and 10% fetal bovine serum (Gemini Biotec). After 72 hours of incubation at 37°C, extensive cytopathic effect (CPE) and large areas of cell clearing/detachment were noted. Flasks were then flash frozen at - 70°C for 8 minutes, thawed briefly at room temperature and the cells and supernatant were then centrifuged at 1500g for 5 minutes followed by a second centrifugation step at 4000g for 5 minutes in order to isolate cell-free viral stocks. Supernatant containing the viral stock was divided into 0.5- and 1.0-ml aliquots in 1.5 ml cryotubes (Nalgene) and archived at −70°C. After freezing, a single tube was thawed, and the viral titer established using both bioassay (TCID50) and real-time RT PCR methods (below).

The tissue culture infectious dose-50 (TCID50) was determined using a viral plaquing assay. CRFK cells were grown in a 96-well tissue culture plate (Genesee Scientific) until the CRFK cells achieved approximately 75-85% confluency. Serial 10-fold dilutions were made of FIPV stock and 200 μL samples of each dilution were added to 10-well replicates. At 72 hours post-infection, the cells were fixed with methanol and stained with crystal violet (Sigma-Aldrich). Individual wells were evaluated visually for virus-induced CPE, scored as CPE positive or negative, and the TCID50 was determined based upon the equation log_10_TCID50 = [total# wells CPE positive/# replicates] + 0.5 to reflect infectious virions per milliliter of supernatant[68].

### Quantification of FIPV by qRT-PCR

Cell-free viral RNA was isolated from the viral stock using the QIAamp Viral RNA Mini Kit (Qiagen) following the manufacturer’s instructions. The isolated RNA was DNase treated (Turbo DNase, Invitrogen) and subsequently reverse transcribed using the High-Capacity RNA-to-cDNA Kit (Applied Biosystems) following the manufacturers’ protocols. The copy number of FIPV and feline GAPDH cDNA were determined using Applied Biosystems’ QuantStudio 3 Real-Time PCR System and PowerUp SYBR Green Master Mix, following the manufacturer’s protocol for a 10 μL reaction. Each PCR reaction was performed in triplicate with water template as a negative control and plasmid DNA as a positive control. A control reaction excluding reverse transcriptase was included in each real-time PCR assay set. cDNA templates were amplified using the FIPV forward primer, 5’-GGAAGTTTAGATTTGATTTGGCAATGCTAG, and the FIP reverse primer, 5’-AACAATCACTAGATCCAGACGTTAGCT (terminal portion of the FIPV 7b gene)[25]. Real-time PCR for the feline GAPDH housekeeping gene was performed concurrently using the primers, 5 GAPDH, 5’-AAATTCCACGGCACAGTCAAG, and 3 GAPDH, 5’-TGATGGGCTTTCCATTGATGA. Cycling conditions for both FIPV and GAPDH amplicons were as follows: 50°C for 2 min, 95°C for 2 min, followed by 40 cycles of 95°C for 15 s, 58°C for 30 s, 72°C for 1 min. The final step included a dissociation curve to evaluate specificity of primer binding. FIPV and GAPDH copy number was calculated based on standard curves generated in our laboratory. Copies of FIPV cDNA determined via real-time RT PCR were normalized per 10^6^ copies of feline GAPDH cDNA.

### Development of anti-helicase chemical fragments

In general, the drugs examined and described in this study were preexisting antiviral agents. In contrast, the helicase enzyme of FIPV was cloned, expressed and utilized as a target for coronavirus and enzyme specific viral discovery. Target DNA sequence of AviTag-FIP Helicase-HisTag was optimized and synthesized. The synthesized sequence was cloned (Adeyemi Adedeji) into vector pET30a with Avi-His tag for protein expression in E. coli. E. coli strain BL21(DE3) was transformed with recombinant plasmid. A single colony was inoculated into 1 L of auto-induced medium containing antibiotic and culture was incubated at 37C at 200rpm.

When the OD600 reached about 3, cell culture was temperature was changed to 15C for 16 hours. Cells were harvested by centrifugation. Cell pellets were resuspended with lysis buffer followed by sonication. The precipitate after centrifugation was dissolved using denaturing agent. Target protein was obtained by one-step purification using Ni column. Target protein was sterilized by 0.22um filter. Yield was 7.2 mg at 0.90 mg/mL, and was stored in PBS, 10% glycerol, 0.5mM L-arginine, pH 7.4. The concentration was determined by Bradford protein assay with BSA as standard. The protein purity and molecular weight were determined by SDS-PAGE with Western blot confirmation.

Surface plasmon resonance (SPR) fragment screening was performed on a ForteBio Pioneer FE SPR platform. A HisCap sensor chip, which contains a NTA surface matrix was used. Channels 1 and 3 were charged with 100uM NiCl2, followed by injection of 50ug/mL FIP protein. Channel 2 was left free of protein, as well as NiCl2, as a reference. Channel 1 was immobilized to a density of ~8000 RU, while channel 3 contained about 12,000 RU. Channel 1 was used. The buffer used for immobilization was 10mM HEPES, pH 7.4, 150mM NaCl, and 0.1% Tween-20. For the assay, DMSO was added to a final concentration of 4%. The proprietary compound library was diluted into the same buffer without DMSO, to a final DMSO concentration of 4% DMSO. Library compounds were screened at a concentration of 100uM using the OneStep gradient injection method. Hits were selected based upon RU and kinetics and utilized for cell-based screening.

### Viral plaquing assay

In order to screen compounds for antiviral activity, infected CRFK cells were treated with compounds in six well replicates and compared to positive control wells (infected cells), negative controls (uninfected cells) and treatment controls (infected cells treated with a known effective antiviral compound) run concurrently on each tissue culture plate. CRFK cells were grown in 96-well tissue culture plates (Genesee Scientific) containing 200 μL culture media. At ~75-85% cell confluency, the media in the uninfected control wells was aspirated and replaced with 200 μL of fresh media. The media in the infected wells was aspirated and replaced with media inoculated with FIPV at a multiplicity of infection (MOI) of 0.004 infectious virions per cell. The tissue culture plate was incubated for 1 hour with periodic gentle agitation (“figure eight” manipulations) performed every 15 minutes to facilitate virus-cell interaction. At 1-hour post-infection, each putative antiviral compound was added to six FIPV-infected wells (to assess compound antiviral efficacy) and six uninfected control wells (to screen for compound cytotoxicity in CRFK cells). All compounds were initially screened at 10 μM, except for the “chemical fragment” compounds supplied by M. Olsen (Midwestern University) which were assessed at 50 μM. The tissue culture plates were incubated for 72 hours at 37°C and subsequently fixed with methanol and stained with crystal violet. Plates were scanned for absorbance at 620 nm using an ELISA plate reader (FilterMax F3, Molecular Devices; Softmax Pro, Molecular Devices). The individual well absorbance values along with the average absorbance value and standard error of the mean for the 6 well experimental replicates were recorded for each treatment condition.

For agents that demonstrated antiviral efficacy in the initial screening at 10 or 50 μM (protected from virus-associated CPE), the EC50 was determined by performing a progressive 2-fold compound dilution series in the viral plaquing assay. For EC50 determination, CRFK cells were grown in 96-well tissue culture plates similarly to that performed for the antiviral screening assay. Aside from the uninfected control wells, all remaining wells were infected with the FIPV as described above. The 2-fold dilution series ranged from 20 μM down to 0 μM and each concentration performed in six well replicates. The number of dilution steps ranged from 6 to 14 and was compound dependent. Six well replicates of uninfected CRFK cells served as a control for normal CRFK cells; six well replicates of CRFK cells infected with FIPV served as untreated, FIPV-infected control wells; and six well replicates of FIPV-infected CRFK cells treated with GS-441524 served as control wells for protection against virus-induced cell death based on published data regarding the efficacy of GS-441524 use *in vitro* in CRFK cells[26].

Tissue culture plates were incubated for 72 hours and subsequently fixed with methanol, stained with crystal violet and scanned for absorbance at 620 nm using an ELISA plate reader. The individual absorbance values along with the average absorbance value and standard deviation for the 6 well experimental replicates were recorded for each treatment condition. The EC50 was calculated by plotting a nonlinear regression equation (dose-response curve) using the program Prism 8 (GraphPad).

### Viral RNA knock-down assay

Real time RT-PCR assays were used to quantify compound inhibition of viral RNA production. CRFK cells were cultured in a 6-well tissue culture plate (Genesee Biotek). At approximately 75-85% cellular confluency, the culture media was replaced with fresh media and the cells were infected with FIPV serotype II at a MOI of 0.2 (MOI based upon the TCID50 bioassay/pfu). Plates were incubated for one hour, with periodic gentle agitation every 15 minutes. FIPV-infected wells were treated with one (monotherapy), two or three (combined anticoronaviral therapy) antiviral compounds; each experimental treatment was performed in triplicate. Compound dosage was based upon the compounds’ EC50 and ranged from 0.001-20 μM. For each experimental set, three culture wells with FIPV-infected and untreated CRFK cells acted as virus-infected controls. The infected cell cultures were subsequently incubated for 24 hours and cell-associated total RNA was isolated using the PureLink™ RNA mini kit (Invitrogen). The RNA was treated with DNAse (TurboDNAse, Ambion), reverse transcribed to cDNA using the High-Capacity RNA-to-cDNA Kit (Applied Biosystems) and FIPV cDNA and feline GAPDH cDNA were measured using real time qRT-PCR, as described above. Fold reduction in viral titer was determined by dividing the normalized average FIPV RNA copy number for untreated, FIPV-infected CRFK cells, into the normalized average FIPV RNA copy number for treated CRFK cells with the compound(s) of interest. The expected additive effect was determined by adding the fold reduction for each monotherapy treatment used in combination. Fold over additive effect was determined by dividing the predicted additive effect into the combined fold reduction value for the particular combined therapy of interest.

### Determination of Cytotoxicity Safety Profiles (CSP)

Compound cytotoxicity in feline cells was assessed using the commercially available kit (CellTox Green Cytotoxicity Assay, Promega) according to the manufacturer’s instructions. Untreated CRFK cells were used as negative controls and cells treated with a cytotoxic solution provided by the manufacturer was used as the positive toxicity control. Briefly, in addition to the control wells, CRFK cells were plated in 96-well tissue culture plates (Genesee Scientific) in four well replicates with 5, 10, 25, 50, or 100 μM concentrations of the compound of interest and were incubated for 72 hours. After 72 hours, the kit DNA binding dye was applied to all wells, incubated at 37°C shielded from light for 15 minutes and the fluorescence intensity at 485-500nm_Ex_/520-530nm_Em_, was subsequently determined using a plate reader (FilterMax F3, Molecular Devices; Softmax Pro, Molecular Devices). Compound cytotoxicity at a particular concentration was assumed to be proportional to the intensity of fluorescence based on the selective penetration and binding of the dye to the DNA of degenerate, apoptotic or necrotic cells. The cytotoxicity range was determined by setting the fluorescence value for cells treated with the positive control reagent as 100% and the untreated feline cells as 0% cytotoxicity. The mean fluorescence value for the four wells containing each compound concentration were then interpolated as a percentage (percent cytotoxicity) ranging from 0-100%.

## Supporting information

**S1 Table.**
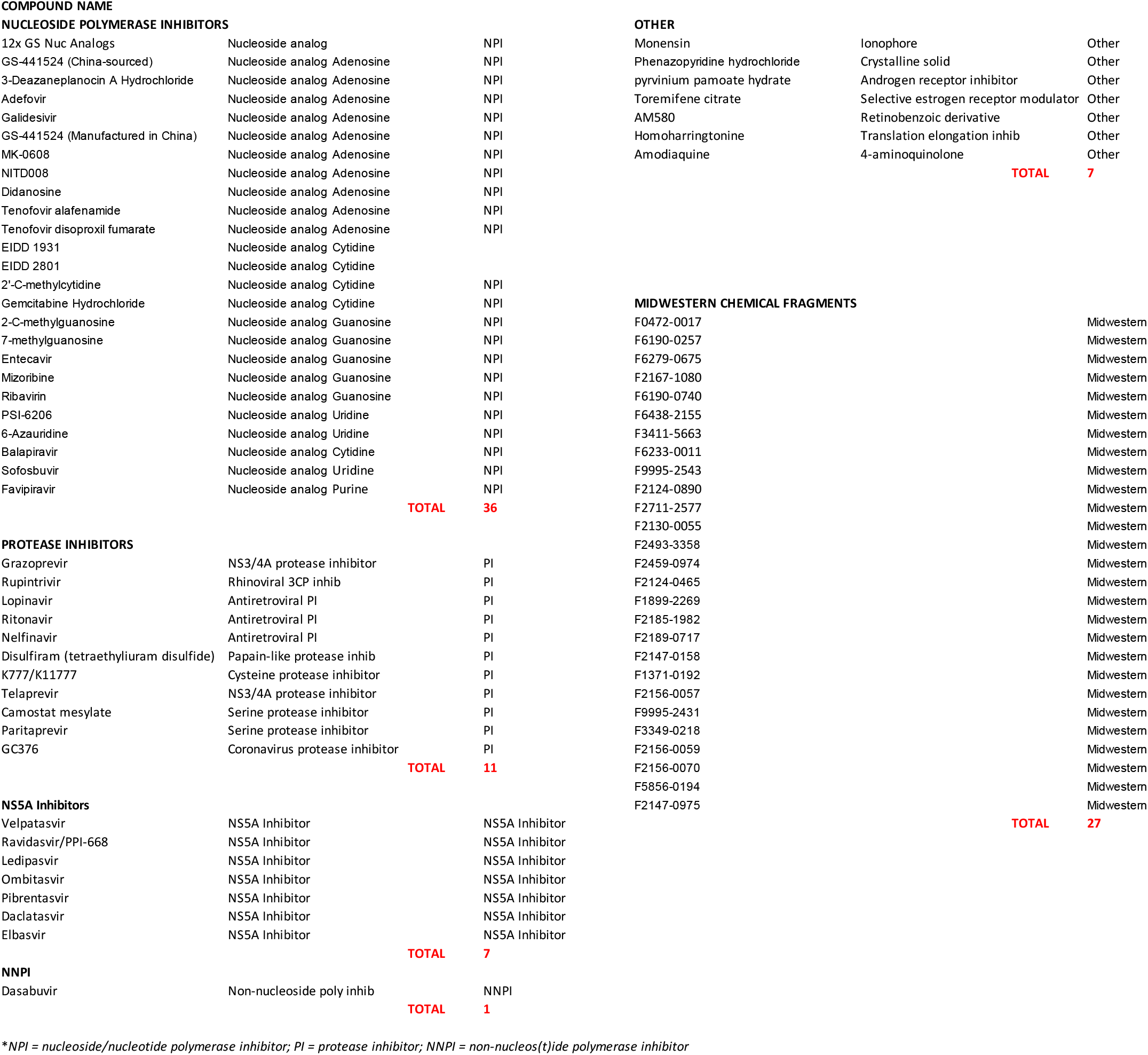
Complete list of compounds screened against FIPV *in vitro*.

NUCLEOSIDE POLYMERASE INHIBITORS
|  |  |  |
| --- | --- | --- |
| 12x GS Nuc Analogs | Nucleoside analog | NPI |
| GS-441524 (China-sourced) | Nucleoside analog Adenosine | NPI |
| 3-Deazaneplanocin A Hydrochloride | Nucleoside analog Adenosine | NPI |
| Adefovir | Nucleoside analog Adenosine | NPI |
| Galidesivir | Nucleoside analog Adenosine | NPI |
| GS-441524 (Manufactured in China) | Nucleoside analog Adenosine | NPI |
| MK-0608 | Nucleoside analog Adenosine | NPI |
| NITD008 | Nucleoside analog Adenosine | NPI |
| Didanosine | Nucleoside analog Adenosine | NPI |
| Tenofovir alafenamide | Nucleoside analog Adenosine | NPI |
| Tenofovir disoproxil fumarate | Nucleoside analog Adenosine | NPI |
| EIDD 1931 | Nucleoside analog Cytidine |  |
| EIDD 2801 | Nucleoside analog Cytidine |  |
| 2'-C-methylcytidine | Nucleoside analog Cytidine | NPI |
| Gemcitabine Hydrochloride | Nucleoside analog Cytidine | NPI |
| 2-C-methylguanosine | Nucleoside analog Guanosine | NPI |
| 7-methylguanosine | Nucleoside analog Guanosine | NPI |
| Entecavir | Nucleoside analog Guanosine | NPI |
| Mizoribine | Nucleoside analog Guanosine | NPI |
| Ribavirin | Nucleoside analog Guanosine | NPI |
| PSI-6206 | Nucleoside analog Uridine | NPI |
| 6-Azauridine | Nucleoside analog Uridine | NPI |
| Balapiravir | Nucleoside analog Cytidine | NPI |
| Sofosbuvir | Nucleoside analog Uridine | NPI |
| Favipiravir | Nucleoside analog Purine | NPI |
|  |  | **TOTAL 36** |
| PROTEASE INHIBITORS |  |  |
| Grazoprevir | NS3/4A protease inhibitor | PI |
| Rupintrivir | Rhinoviral 3CP inhib | PI |
| Lopinavir | Antiretroviral PI | PI |
| Ritonavir | Antiretroviral PI | PI |
| Nelfinavir | Antiretroviral PI | PI |
| Disulfiram (tetraethyliuram disulfide) | Papain-like protease inhib | PI |
| K777/K11777 | Cysteine protease inhibitor | PI |
| Telaprevir | NS3/4A protease inhibitor | PI |
| Camostat mesylate | Serine protease inhibitor | PI |
| Paritaprevir | Serine protease inhibitor | PI |
| GC376 | Coronavirus protease inhibitor | PI |
|  |  | **TOTAL 11** |
| NS5A Inhibitors |  |  |
| Velpatasvir | NS5A Inhibitor | NS5A Inhibitor |
| Ravidasvir/PPI-668 | NS5A Inhibitor | NS5A Inhibitor |
| Ledipasvir | NS5A Inhibitor | NS5A Inhibitor |
| Ombitasvir | NS5A Inhibitor | NS5A Inhibitor |
| Pibrentasvir | NS5A Inhibitor | NS5A Inhibitor |
| Daclatasvir | NS5A Inhibitor | NS5A Inhibitor |
| Elbasvir | NS5A Inhibitor | NS5A Inhibitor |
|  |  | **TOTAL 7** |
| NNPI |  |  |
| Dasabuvir | Non-nucleoside poly inhib | NNPI |
|  |  | **TOTAL 1** |
| OTHER |  |  |
| Monensin | Ionophore | Other |
| Phenazopyridine hydrochloride | Crystalline solid | Other |
| pyrvinium pamoate hydrate | Androgen receptor inhibitor | Other |
| Toremifene citrate | Selective estrogen receptor modulator | Other |
| AM580 | Retinobenzoic derivative | Other |
| Homoharringtonine | Translation elongation inhib | Other |
| Amodiaquine | 4-aminoquinolone | Other |
|  | **TOTAL** | **7** |

MIDWESTERN CHEMICAL FRAGMENTS
|  |  |
| --- | --- |
| F0472-0017 | Midwestern |
| F6190-0257 | Midwestern |
| F6279-0675 | Midwestern |
| F2167-1080 | Midwestern |
| F6190-0740 | Midwestern |
| F6438-2155 | Midwestern |
| F3411-5663 | Midwestern |
| F6233-0011 | Midwestern |
| F9995-2543 | Midwestern |
| F2124-0890 | Midwestern |
| F2711-2577 | Midwestern |
| F2130-0055 | Midwestern |
| F2493-3358 | Midwestern |
| F2459-0974 | Midwestern |
| F2124-0465 | Midwestern |
| F1899-2269 | Midwestern |
| F2185-1982 | Midwestern |
| F2189-0717 | Midwestern |
| F2147-0158 | Midwestern |
| F1371-0192 | Midwestern |
| F2156-0057 | Midwestern |
| F9995-2431 | Midwestern |
| F3349-0218 | Midwestern |
| F2156-0059 | Midwestern |
| F2156-0070 | Midwestern |
| F5856-0194 | Midwestern |
| F2147-0975 | Midwestern |
|  | **TOTAL 27** |

